# The *Drosophila tyramine-beta-hydroxylase* gene encodes multiple isoforms with different functions

**DOI:** 10.1101/2024.06.10.598396

**Authors:** Manuela Ruppert, Stefanie Hampel, Nagraj Sambrani, Osman Cibik, Gerbera Classen, Andrea Duenisch, Claire Fuchs, Thomas Kell, Sravya Paluri, Henrike Scholz

## Abstract

The Tyramine-beta−hydroxylase (Tbh) is required for octopamine synthesis. To better understand the function of Tbh in neurotransmitter synthesis, we analyzed the molecular genetic organization of the *Drosophila melanogaster Tbh* gene and found that the *Tbh* gene encodes multiple transcripts. The transcripts differ in their 5’UTR, which results in proteins that differ in their size and putative phosphorylation sites, suggesting that the *Tbh* function is regulated at translational and posttranslational levels. We generated a new *Tbh* mutant – *Tbh^Del3^* - using FLP/FRT recombination mutagenesis to remove the translational start site still that is present in *Tbh^nM18^*mutants. The *Tbh^Del3^* mutants share ethanol tolerance and larval locomotion defects with the *Tbh^nM18^* mutants. But, they differ in terms of their cellular stress response. To develop normal levels of ethanol tolerance, Tbh is required in a subset of Tbh expressing neurons in the adult brain, which was identified using a newly generated *Tbh*-Gal4 driver. Taking advantage of a newly generated Tbh antibody serum, we show that one Tbh isoform is expressed in a group of peptidergic Hugin-positive and noradrenergic neurons uncoupling Tbh function from octopamine synthesis. The existence of different functional Tbh isoforms impacts our understanding of the regulatory mechanisms of neurotransmitter synthesis and the function of the octopaminergic neurotransmitter system in cellular processes and the regulation of behavior.

**Author Summary:** Vertebrates and insects have structurally identical signaling molecules in their nervous system, such as the neurotransmitter dopamine. But, there are also neurotransmitters that are thought to only occur in the vertebrate or insect brain. Noradrenaline is one such neurotransmitter that regulates flight and fight responses in vertebrates. In insects such as the fruit fly *Drosophila melanogaster*, the structurally very similar neurotransmitter octopamine is considered to be an invertebrate-specific neurotransmitter that performs similar functions to noradrenaline. The functional similarities also extend to enzymes required for synthesis. Our analysis shows that the enzyme for octopamine synthesis exists in several variations and that the connection between the enzymes and the synthesized neurotransmitter may not be as simple as presumed. Exploiting molecular, behavioral and neuroanatomical studies, we show that different variants might be used in response to different environmental conditions and/or the synthesis of alternative, structurally similar neurotransmitters, such as noradrenaline. These results challenge our view on the functions of octopamine and noradrenaline in the regulation of behavior.

## Introduction

The octopaminergic neurotransmitter system regulates physiological processes ranging from energy metabolism to behavior. The rate-limiting enzyme in *Drosophila melanogaster* required for octopamine synthesis is the Tyramine-beta-hydroxylase (Tbh) enzyme that efficiently hydrolyzes tyramine to octopamine (1) (2). The originally identified Tbh isoform shares approximately 39% identity with the mammalian Dopamine-beta*-* hydroxylase (Dbh; (1)). In vertebrates, Dbh catalyzes the hydroxylation of dopamine to noradrenaline, a molecule structurally very similar to octopamine (3).

To analyze the function of octopamine in regulating physiological processes in *Drosophila melanogaster,* mutants were generated using P-element excision mutagenesis (1). One of these mutants, the *Tbh^nM18^* mutant has no detectable level of octopamine and an approximately tenfold increase in the tyramine levels (1). Initial phenotypic characterization of this mutant revealed that Tbh is required for female fertility, a function to which a number of octopaminergic neurons in the thoracic-abdominal ganglion of the adult female fly contribute (1); (4). The mutant is now intensively used to study the function of Tbh and/or octopamine in the regulating cellular processes and behavior during development or in the adult nervous system.

Tbh is required to initiate and adjust behaviors, yet does not appears to be directly involved in the performance of motor behavior. For example, adult *Tbh^nM18^* flies can run as fast as the controls, but they show reduced initiation of locomotion - such as aggression or flight activity (5); (6), (7). In addition, they show a reduced likelihood of extending their proboscis in response to sucrose applied to their tarsi, however, when responding, they habituate just as well as the control flies (8).

A strong activator of locomotor activity is ethanol (9); (10); (11). Brief exposure of the odorant ethanol induces an olfactory startle response, measured by increased locomotor activity (10), (5). The startle response is strongly reduced in *Tbh^nM18^* mutants and is due to loss of octopamine function (5). In contrast to the initial reduction in locomotor activity, prolonged ethanol exposure leads to increased locomotor activity in adult *Tbh^nM18^* mutants (5). Moreover, our previous work shows that approaching an attractive stimulus, such as ethanol, requires the function of Tbh, given that octopamine is essential in a subset of Tbh-positive neurons (13); (12). Compared to control flies, *Tbh^nM18^* mutant flies make random choices in a choice essay, which as well supports the hypothesis that the initiation requires (12). These results further show that the execution of motor output is not altered in adult *Tbh^nM18^* mutants, but that there is a lack of initiation to perform the required action.

In addition to a reduced initial behavioral response rate, *Tbh^nM18^* flies fail to adapt their behavior after a previous experience. The *Tbh^nM18^* flies show normal sensitivity to the effect of ethanol on postural control, but are unable to increase their resistance to a similar extend as controls and, therefore, develop reduced ethanol tolerance (14). After multiple exposures to ethanol, they build up ethanol tolerance and eventually reach the same levels as controls (10). Accordingly, after chronic, long-term exposure to ethanol their tolerance does not differ from control flies (15).

To further examine the function of Tbh in regulating ethanol-induced behaviors, we first analyzed the molecular nature of the *Tbh* gene using RT-PCR analysis. Because we identified additional splice variants of *Tbh* and still detected residual transcripts in the *Tbh^nM18^* mutants, we generated a new *Tbh* allele using FLP/FRT recombination-based mutagenesis. We tested this new mutant for its ability to develop ethanol tolerance and to respond to cellular stress. To uncover differences in the regulation of locomotion in *Tbh* mutants, we characterized adult and larval locomotion in response to external stressors and changes of internal motivation. We next determined under what circumstances *Tbh* is required in the nervous system for tolerance development. By restoring Tbh expression in *Tbh^nM18^* mutants using a newly generated *Tbh*-promoter-Gal4 driver, we identified neurons required for tolerance. Employing a newly generated Tbh antibody, we confirmed that these neurons do indeed express Tbh. To analyze whether different Tbh isoforms are expressed in the same cells, we analyzed the Tbh expression pattern using different Tbh antibodies. Our results are significant, because the generation and phenotypic characterization of a new *Tbh* mutant along with the identification of multiple Tbh isoforms with different functions, provide new insights into the regulation of neurotransmitter synthesis and the regulation of behavior.

## Results

The *Tbh^nM18^* mutant, which lacks the neurotransmitter octopamine, is often used to study the function of Tbh and/or octopamine. This mutant was isolated by P-element excision mutagenesis using the MF372 insertion line as starting point. However, the exact molecular change responsible for the mutation has not been described in detail (1).

### The *Tbh* gene encodes several transcripts

First, we determined the precise nature of the molecular lesion. Through sequencing analysis, we found that the *Tbh^nM18^* allele carries a deletion of 4269 bp, including 32 bp of the coding sequence of the 3’ end of the second exon and sequence of the second intron (Fig 1A). The deletion does not include the translation start site of the *Tbh* gene, but results in a frame shift of the coding sequence. To analyze the consequences of the deletion at transcript level, we determined the amount of *Tbh* transcript in the mutants with exon-specific primers for the transition between exon 5 and 6 (*Tbh^5-6^*) using quantitative real-time PCR (qRT-PCR; Fig 1B). In the *Tbh^nM18^* mutant, the *Tbh^5-6^* transcript sequence is still present and the amount is not significantly reduced. To determine whether additional *Tbh* transcripts exist that may not be affected by the mutation, we analyzed the sequence of EST clones of *Tbh*. Furthermore, we compared the *Tbh* transcripts of *Periplaneta americana* (16) with the genome of *Drosophila melanogaster.* In addition, we analyzed the sequence of the second intron for additional open reading frames. To validate the transcripts, we performed RT-PCR analysis on cDNA isolated from heads of control flies. We identified four different transcripts and a novel exon within the second intron of the annotated *Tbh* gene (*Tbh^RA^* to *Tbh^RD^*; Fig 1C; for sequence of transcripts S1 Text). The *Tbh^RA^* and *Tbh^RB^* transcripts differ in length of the 5’ UTR, but not in the protein they encode. In the *Tbh^RC^* transcript, the second exon is alternatively spliced. The fourth 1.5 kb-long *Tbh^RD^* transcript contains the sequence of a newly identified exon located after the second annotated exon. In our RT-PCR analysis we did not detect any connections between the new third exon and the first and/or second exon of *Tbh^RA/RB^*.

**Figure 1:**
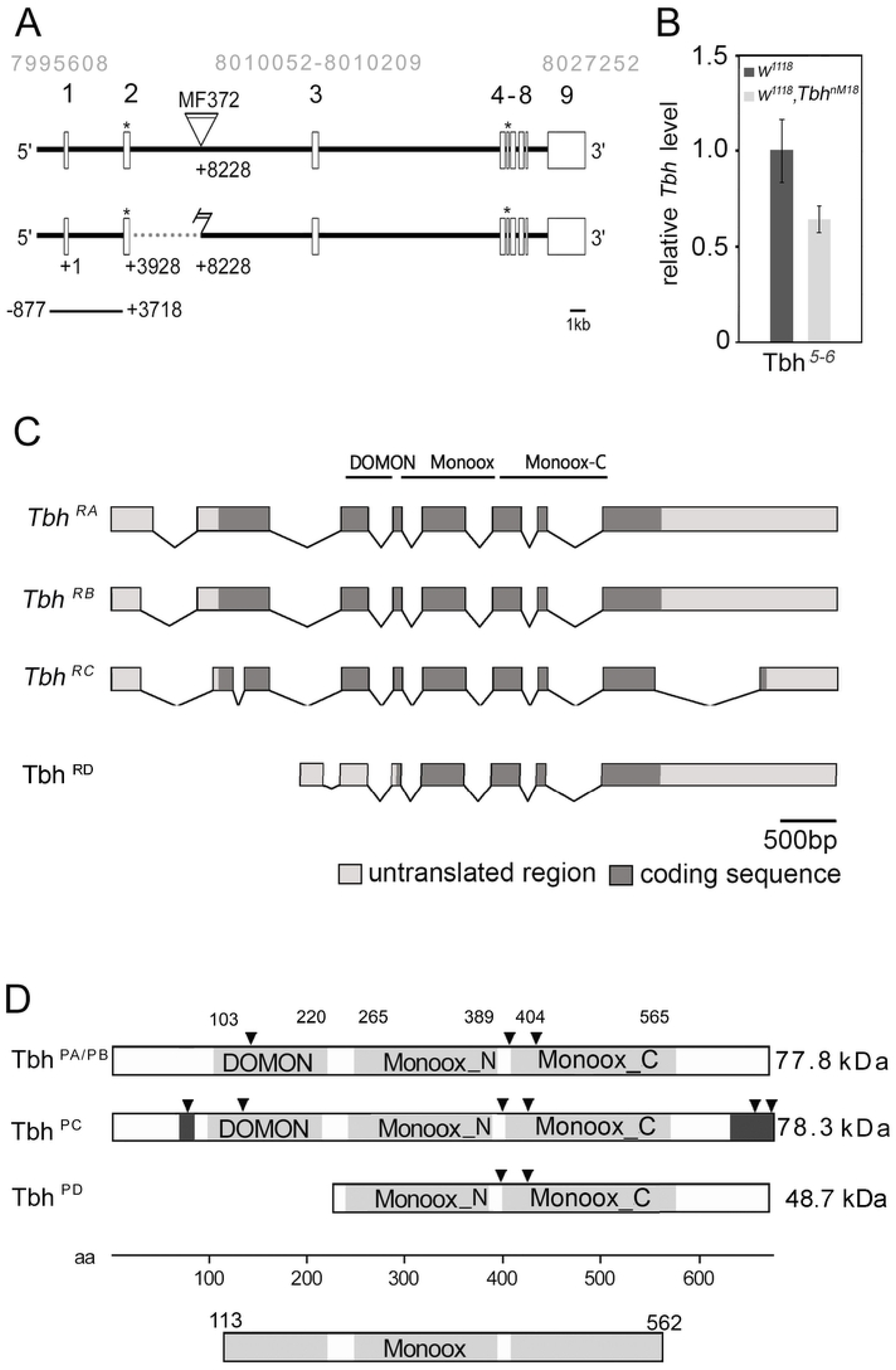
The *Tbh* gene encodes several isoforms. The genomic organization of the *Tbh* gene spans 31 kb. The triangle indicates the transposable element of the *Tbh^MF372^* insertion, the light gray numbers describe the genomic region of the *Tbh* gene and the position of the newly identified third exon. The dotted line represents the lesion of the mutant *Tbh^nM18^* allele. The deletion includes 32 bp of the second exon, but does not include the translational start sites marked with the asterisks. The bar below indicates the genomic region used to generate the *4.6-Tbh-*Gal4 driver. (B) The qRT-PCR analysis of poly(A)-selected RNA form *Tbh^nM18^* heads using *Tbh* primers specific to the transition between exon 5 to 6 shows *Tbh* transcripts in the mutants (N = 4 independent cDNA samples with three replicates; *RpLPO* gene primers were used as reference; the transcript levels were not different as determined by Student’s *t*-test. Error bars are s.e.m.). (C) Four *Tbh* splice variants were identified using RT-PCR analysis. Three transcripts had common sequences for the DOMON domain and the two copper-type II ascorbate-dependent monooxygenase domains. (D) Tbh^PA/PB^ and Tbh^PC^ share the DOMON domain and two copper type II ascorbate-dependent monooxygenase domains (light gray boxes). The Tbh^PC^ protein has unique amino acid sequence at the N- and C-terminus (dark gray boxes). The Tbh^PD^ protein lacks the DOMON domain. The putative protein kinase C phosphorylation sites are marked with black triangles. The sequence of transcripts and proteins is found in S1 Text.

The four transcripts encode three different proteins, because the *Tbh^RA^* and *Tbh^RB^* transcripts encode the same protein. The Tbh^PA/PB^ and Tbh^PC^ isoforms of 77,7 kDA and 78,3 kDA, respectively, share the DOMON domain and the N- and C-terminus of the copper type II ascorbate-dependent monooxygenase domain (Fig 1D; for sequence of proteins S1 Text). The DOMON domain is a 110 to 125 amino acid stretch in the N-terminal region of the dopamine beta-monooxygenase that is found in combination with monooxygenase domains (17). The Tbh^PC^ isoform differs from Tbh^PA/PB^ in its N- and C-terminal regions, which contain three additional putative protein kinase C phosphorylation sites (Fig 1D). The Tbh^RD^ protein is approximately 48,7 kDA in size and contains a similar C-terminal region and a truncated N-terminus affecting the DOMON domain. In summary, the *Tbh* gene transcribes at least four different transcripts and encodes at least three different isoforms. Furthermore, the *Tbh^nM18^* mutant allele does not appear to be a transcriptional null allele.

### The *Tbh^Del3^* mutant shows defects in the cellular stress response

Because the *Tbh^nM18^* mutation did not delete the translational start site of *Tbh^RA^* and *Tbh^RB^* and we detected residual *Tbh* transcripts in *Tbh^nM18^* mutants, we decided to generate a new mutant with a missing translation start site using FLP/FRT recombination-based mutagenesis (18). Transheterozygous flies carrying the FRT-site containing transposons *XP^d01344^* upstream and *XP^d10000^* downstream of the second exon of the *Tbh* gene were used as starter lines (Fig 2A). Activation of the FLP recombinase in these flies resulted in the removal of the genomic sequence between the two transposons and fragments of the transposable elements. The newly generated *Tbh^Del3^* allele contains a 9.2 kb deletion including the first and second exons of *Tbh* as confirmed by PCR analysis (Fig 2A).

**Figure 2:**
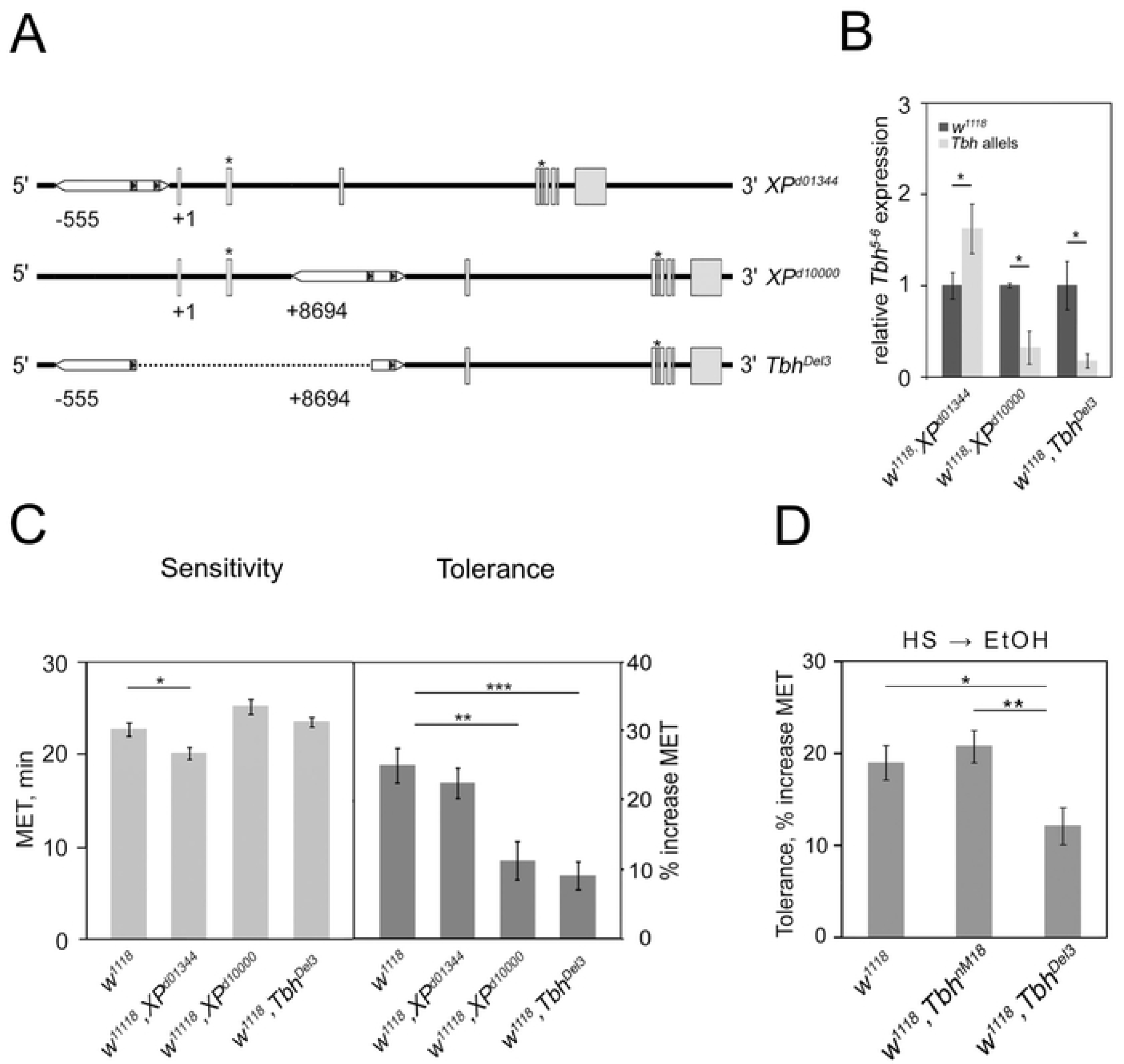
The *Tbh^Del3^* mutant is impaired in the cellular stress response. (A) The genomic organization of the *XP^d01344^* and the *XP^d1000^* lines used to generate the *Tbh^Del3^* mutant is shown. The dotted line indicates the deletion of the *Tbh^Del3^* allele generated by FRT/Flip-based recombination. (B) The qRT-PCR analysis of *XP^d01344^ XP^d1000^* lines and *Tbh^Del3^* mutant revealed significantly up- or down-regulation of *Tbh* transcripts (N = 4 independent cDNA samples with three replicates; *RpLPO* gene primers were used as reference; Student’s *T*-test \**P* < 0.05. Error bars are s.e.m.). (C) The *Tbh^XPd01344^* mutant is significantly more sensitive to ethanol than the controls. The*Tbh^Del3^* and *Tbh^XPd10000^* mutants developed similarly significantly reduced ethanol tolerance (N = 14 – 23). (D) After a heat shock, control and *Tbh^nM18^* are more resistant to ethanol. The *Tbh^Del3^* mutants developed significantly less resistance to ethanol (N = 14 – 17). To calculate the heat-ethanol cross tolerance, the MET of heat-shocked flies is compared to the MET1 of flies of the same genotype without heat shock. Significance was determined with ANOVA, Tukey-Kramer *post hoc* test and for qRT-PCR analysis with Student’s *t*-test. In all panels, asterisk indicate significant values as follows: \**p* < 0.05; \*\**p* < 0.01 and \*\*\**p* < 0.001. The error bars represent s.e.m.. For data see S2 Table.

Next, we used qRT-PCR to analyze whether the new deletion changes the *Tbh* transcript level (Fig 2B). We used the parental lines for comparison. In the *Tbh^Del3^* mutants, the *Tbh^5-6^* transcript fragment level is significantly reduced by about 83% and in the insertion line *XP^d1000^* by 68%. In contrast, in the *XP^d01344^* insertion line, the *Tbh^5-6^* transcript fragment is significantly increased by about 62%. Because the *Tbh* transcript levels were significantly altered in the *XP^d01344^* and *XP^d10000^* lines, the lines are considered *Tbh* mutants.

The *Tbh^nM18^* mutants develop reduced ethanol tolerance (10). To find out whether the *Tbh^Del3^* mutants also have reduced ethanol tolerance, we analyzed the ethanol sensitivity and tolerance of the *Tbh^Del3^* mutants and the parental insertion lines (Fig 2C). To measure ethanol sensitivity and tolerance, flies were inserted into the “inebriometer” – an assay that measures the effect of ethanol on postural control in a population of flies (19). After the first exposure and a second exposure four h later, flies typically show an increased resistance to the effect of ethanol on postural control. This relative increase is defined as ethanol tolerance (10). To quantify the behavior, the average time a population spent in the column is determined as *mean elution time* (MET). The ethanol sensitivity of *Tbh^Del3^* and *Tbh^XPd1000^* did not differ from the controls, but the *Tbh^XPd01344^* flies showed a significantly increased sensitivity of approximately 11% (Fig 2C). The control flies developed ethanol tolerance of approximately 25% similar to *Tbh^XPd01344^* flies. In contrast, the *Tbh^Del3^* mutants significantly reduced their tolerance by around 64% and the *Tbh^XPd10000^* mutants by around 55% (Fig 2C). The *Tbh^Del3^* and the *Tbh^d10000^* mutants exhibit a similar reduced tolerance phenotype as the *Tbh^nM18^* mutants, but fail to complement the female sterility of *Tbh^nM18^* mutants (data not shown).

The *Tbh^nM18^* mutants are more resistant to ethanol when exposed to 30-min heat shock of 37 °C four h before ethanol exposure, similar to control flies (14). To determine whether the new *Tbh^Del3^* mutant also develops heat-induced resistance to ethanol, we analyzed the ethanol-induced behavior after heat shock of *Tbh^Del3^*, *Tbh^nM18^* mutants and controls (Fig 2D). In *Tbh^nM18^* mutants, prior heat exposure resulted in approximately 22% increased resistance to ethanol compared to the ethanol-induced behavior of mutant flies not exposed to heat. This increased in ethanol resistance was comparable to that of control flies. In contrast, the *Tbh^Del3^* mutant flies showed significantly reduced resistance to ethanol by approximately 35% after heat shock (Fig 2D). In conclusion, the new *Tbh^Del3^* mutants, like the *Tbh^nM18^* mutants, exhibit reduced ethanol tolerance but differ in their resistance to ethanol after heat shock.

### The *Tbh* mutants respond to external stimulation and internal motivation with increased locomotion

*Tbh* mutants express normal ethanol sensitivity, which indicates they have the motor skills to perform the task required in a behavioral assay. However, they fail to adjust their behavioral output at a second ethanol exposure. To further investigate the motor performance, we analyzed how well adult and larval *Tbh^nM18^* and *Tbh^Del3^* mutants can move (Fig 3). We filmed the flies for one minute and analyzed their walking trajectories (Fig 3A and Fig 3B). *Tbh^nM18^* flies moved significantly slower than control flies by about 19% and *Tbh^Del3^* by about 28% (Fig 3A). Additionally, *Tbh^nM18^* flies traveled similar distance as the control flies, while *Tbh^Del3^* flies traveled approximately 35% less distance (Fig 3B).

**Figure 3:**
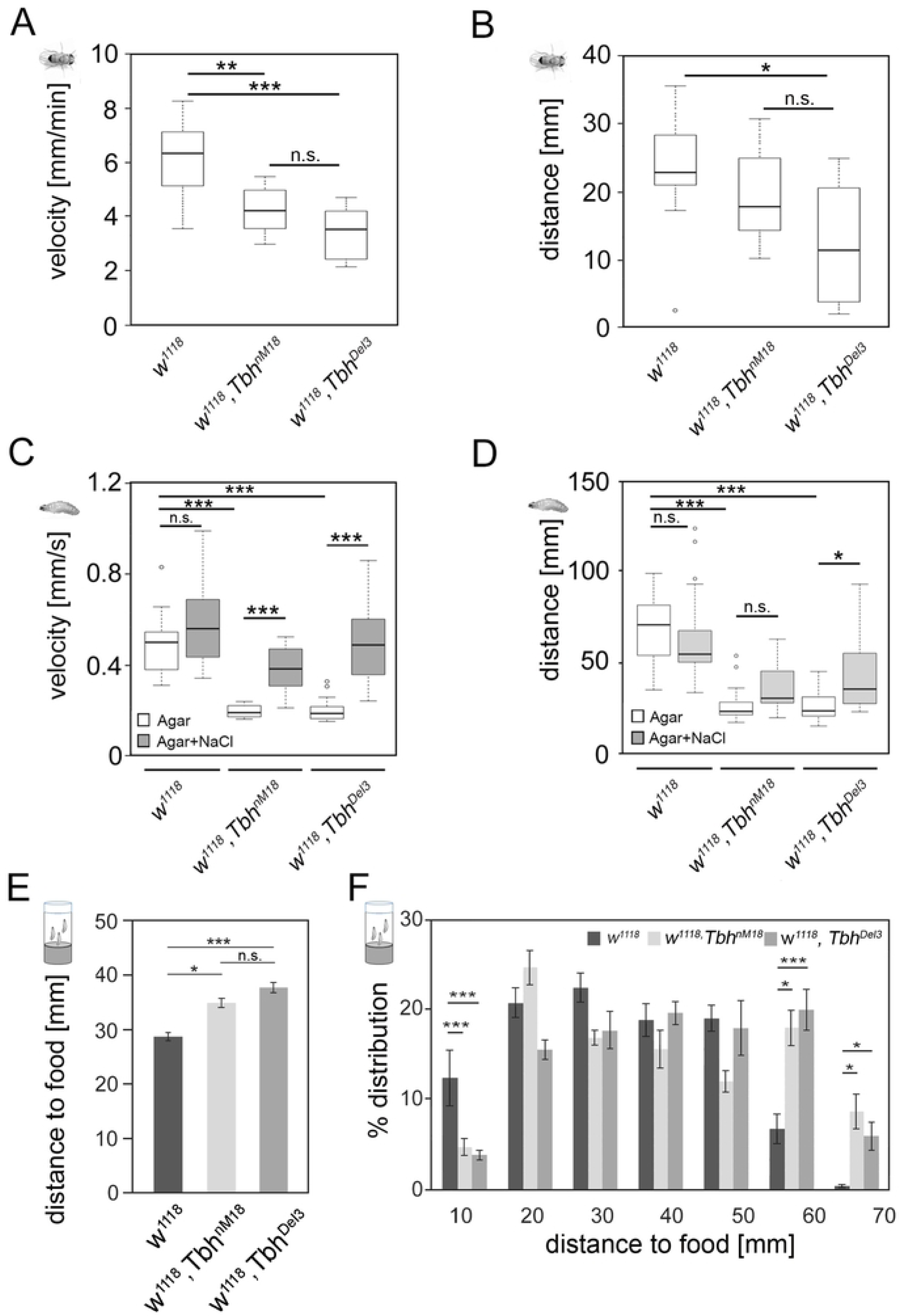
Reduced movement of *Tbh* mutants is improved by external stimulation and internal motivation. The speed (A) and the distance traveled (B) of individual male flies were analyzed in a glass arena. The *Tbh^nM18^* and *Tbh^Del3^* mutants walked significantly slower and *Tbh^Del3^* mutants walked less fat than controls (N = 11 – 13 flies). (C) Both *Tbh* mutant larvae moved significantly slower but increased speed when exposed to salt. (D) Similarly, the *Tbh* mutant larvae traveled a significantly shorter distance, and the *Tbh^Del3^* mutants significantly improved the distance when exposed to 2 M NaCl salt (N = 11 – 13 larvae per genotype). (E) The 3^rd^ instar of *Tbh* mutant larvae moved significantly further out of the food to pupate than controls. (F) Analysis of the distribution pattern of the distances traveled collected in 10 mm bins showed that the *Tbh^nM18^*and *Tbh^Del3^* larvae moved significantly further from the food to pupate (N = 9 - 10 vials with at least 50 pupae per vial). For non-normal distributed data in (A), (C) and (D) the Kruskal-Wallis test followed by a Bonferroni correction were used to determine differences. Normal distributed data were compared using ANOVA Tukey-Kramer *post hoc* analysis. The asterisks indicate significance as follows * *p* < 0.016; *\*\* p* < 0.003 and *\*\*\* p* < 0.0016 for (A) and (B); *\* p* < 0.0083; *\*\* p* < 0.0017 and *\*\*\* p* < 0.00083 for (C) and (D) and * *p* < 0.05; *\*\* p* < 0.01 and *\*\*\* p* < 0.001 for (E) and (F). Error bars are s.e.m. for (E) and (F). For data see S2 Table.

To examine the movement of larvae, control and *Tbh* mutant larvae were placed on agar plates and their movements were monitored for two minutes. Consistent with previous results, *Tbh^nM18^* mutant larvae moved at significantly reduced speed and crawled less far (20); (21) (Fig 3C and Fig 3D). *Tbh^Del3^* mutants behaved similarly (Fig 3C and Fig 3D).

Adult *Tbh^nM18^* mutants increase their locomotion and speed when exposed to ethanol, suggesting that the environmental stimulus and/or the motivation to move is not strong enough to elicit the same locomotor pattern without ethanol (5). To encourage *Tbh^nM18^* mutant larvae to move, we exposed them to an aversive stimulus by adding 2 M NaCl to the agar plate (Fig 3C and Fig 3D). Control larvae did not increase their movement speed and distance traveled significantly. *Tbh^nM18^* mutants increased speed twofold and *Tbh^Del3^* mutants moved 2.5 times as fast and crawled 68% farther.

To analyze whether the intrinsic motivation influences the movement pattern, we measured the distance 3^rd^ instar larvae crawl out of the food to pupate (Fig 3E and Fig 3F). *Tbh^nM18^* and*Tbh^Del3^* mutant larvae crawled 22 – 31% farther out of the food to pupate than the control larvae (Fig 3E). Analysis of the distribution pattern of the distance traveled shows that *Tbh^nM18^*and*Tbh^Del3^* larvae travelled significantly longer distances from food to pupate (Fig 3F). Therefore, 3^rd^ instar larvae of *Tbh* mutant larvae have the ability to cover long distances. Taken together, the observations that *Tbh* mutants develop normal ethanol sensitivity and walk as quickly and at least as long as controls when exposed to salt suggest that *Tbh* mutants do not have defects in their locomotor ability. Rather, they exhibit deficiencies in translating a previous experience into a new behavior and have defects in adapting their behavior in response to changes in the external and internal environment.

### Tbh is required in the adult flies for ethanol tolerance

To determine at which developmental stage Tbh is required for ethanol tolerance, we wanted to restore Tbh expression in the *Tbh^nM18^* mutants at the adult stage using a heat-inducible *Tbh* transgene. First, we analyzed whether the *white* gene, used as marker for the P-element insertion of the transgenes, influences ethanol sensitivity and the development of ethanol tolerance. Therefore, we tested *w*^+^, *Tbh^nM18^* and *w*^-^, *Tbh^nM18^* and their respective controls for ethanol sensitivity and tolerance (Fig 4A). Both *w^+^, Tbh ^nM18^* and *w^1118^, Tbh^nM18^* mutants developed normal ethanol sensitivity and a similar significantly reduced tolerance by approximately 40% to 55%. Thus, the *white* mutation had no effect on the sensitivity or reduced tolerance phenotype of *Tbh* mutants.

**Figure 4:**
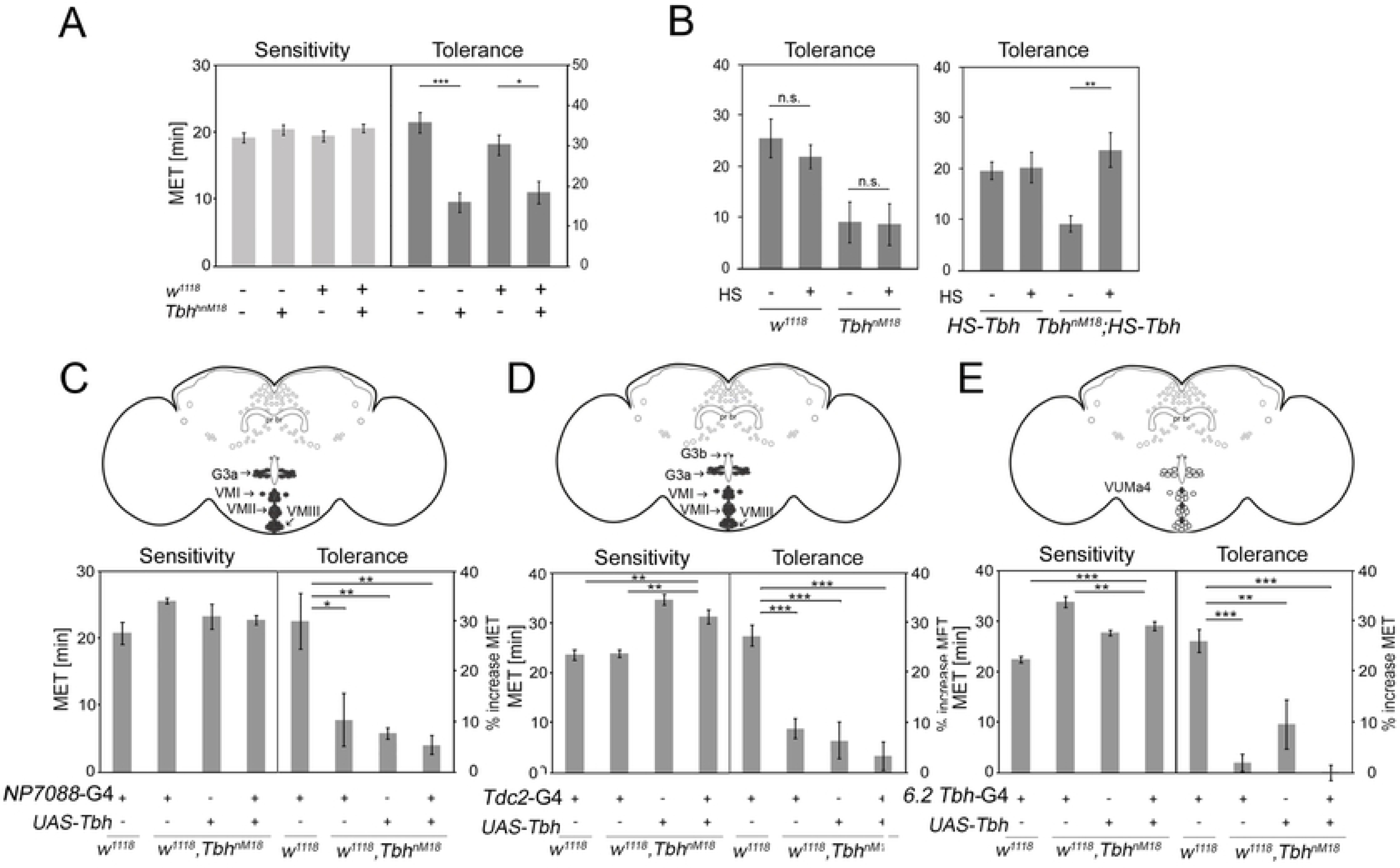
Tbh is required for ethanol tolerance in adult flies. (A) The *w^1118^* mutation did not influence ethanol sensitivity or reduced ethanol tolerance of *Tbh^nM18^* mutants (N = 10 – 16). (B) Controls and *Tbh^nM18^* mutants received a 30-min heat shock of 37° 16 h before the first ethanol exposure. Heat shock had no effect on tolerance development in the controls or the reduced tolerance of *Tbh^nM18^*mutants (N = 13). The reduced tolerance of the *Tbh^nM18^* mutants was restored to normal levels by induction of the *hs-Tbh* transgene (N = 5 – 7). The expression of *Tbh* under the control of (C) the *NP7088*-Gal4 (N = 4), (D) the *dTdc2*-Gal4 (N = 12 - 17) or (E) the *6.2-Tbh-*Gal4 driver (N = 6) did not restore the reduced ethanol tolerance of the *Tbh^nM18^* mutants. In the schemata, Tbh-positive neurons that are addressed by the respective Gal driver are shown in gray filled circles. The ANOVA Tukey-Kramer *post-hoc* test was used to determine difference between groups. Non-significant differences are reported indicated as n.s. and the asterisks indicate \**p* < 0.05 and \*\**p* < 0.01. Error bars are s.e.m.. For data see S2 Table.

We have previously shown that a heat shock of 37 °C four h before ethanol exposure increased the resistance to ethanol (14). To avoid the possible influence of the 37 °C heat shock on ethanol tolerance, the heat shock was administered 16 h before the first ethanol exposure. As additional control, we analyzed the effect of heat shock treatment on the development of ethanol tolerance in controls and *Tbh^nM18^* mutants (Fig 4B, left). Heat treatment 16 h before ethanol exposure had no effect on the tolerance of controls and the reduced tolerance of the *Tbh^nM18^*mutants. Next, we used the same procedure to induce the *hs*-*Tbh* transgene in the *Tbh^nM18^* mutants (Fig 4B, right). In controls, the heat shock had no effect on tolerance, but the reduced tolerance phenotype of the *Tbh^nM18^*mutants was significantly improved to control levels (Fig 4B, right). This led us to conclude that Tbh is required for ethanol tolerance in the adult stage.

To identify in which neurons *Tbh* is required for tolerance, we restored Tbh expression in the *Tbh^nM18^* mutants using different Gal4 drivers. The Gal4 lines drive transgene expression in subsets of Tbh-positive neurons (Fig 4C to Fig 4E; (13)). Tbh expression under the control of the tyraminergic/octopaminergic *dTdc2*-Gal4 and *NP7088*-Gal4 driver lines (22), and the *6.2-Tbh*-Gal4 driver (13) did not significantly improve the reduced ethanol tolerance of *Tbh^nM18^* mutants (Fig 4C to Fig 4E). Thus, none of the Gal4-targeted Tbh-positive neurons, including the ventral unpaired median neurons (VUMs), the G3a, or the G3b neurons, appeared to regulate Tbh-dependent ethanol tolerance.

### A subset of neurons mediates Tbh-dependent ethanol tolerance

Because the *dTdc2*-Gal4 and 6.2-*Tbh*-Gal4 drivers do not target the entire set of Tbh-positive neurons (13), we generated a new *Tbh-*promoter-Gal4 driver. For this, we used a 4.6kb fragment spanning the genomic region -877 to +3718 of the *Tbh* gene as a promoter fragment (indicated in Fig 1A) and characterized the expression pattern of the *4.6-Tbh*-Gal4 driver in the brain of adult flies using the *UAS-mcD8::GFP* transgene (Fig 5A). GFP expression was detected in approximately 61 - 72 cells (Fig 5A to Fig 5A′′). In each hemisphere, the following clusters were identified: a cluster of 11 to 13 cells innervating the ellipsoid body (EB), a cluster of 16 to 18 cells innervating the antennal lobes (AL), and one cell localized in the pars intercerebralis (PI) projects towards the esophagus. In addition, there are two clusters in the posterior part of the brain: one cluster with two cells (PC1) and a second cluster with two to three cells (PC2). Strong Johnston’s Organ expression was also found projecting into the antennal-mechanosensory and motor center (AMMC) (Fig 5A′). Expression of *UAS-Tbh* under the control of the *4.6-Tbh-*Gal4 driver restored the reduced ethanol tolerance of the *Tbh^nM18^* to control levels (Fig 5B). To determine whether the dose of Tbh enzyme is important for normal ethanol tolerance development, we overexpressed Tbh in control flies with the 4.6-*Tbh*-Gal4 driver (Fig 5C). The increased levels of *Tbh* significantly reduced ethanol tolerance, suggesting that Tbh level needs to be tightly regulated for tolerance to develop.

**Figure 5:**
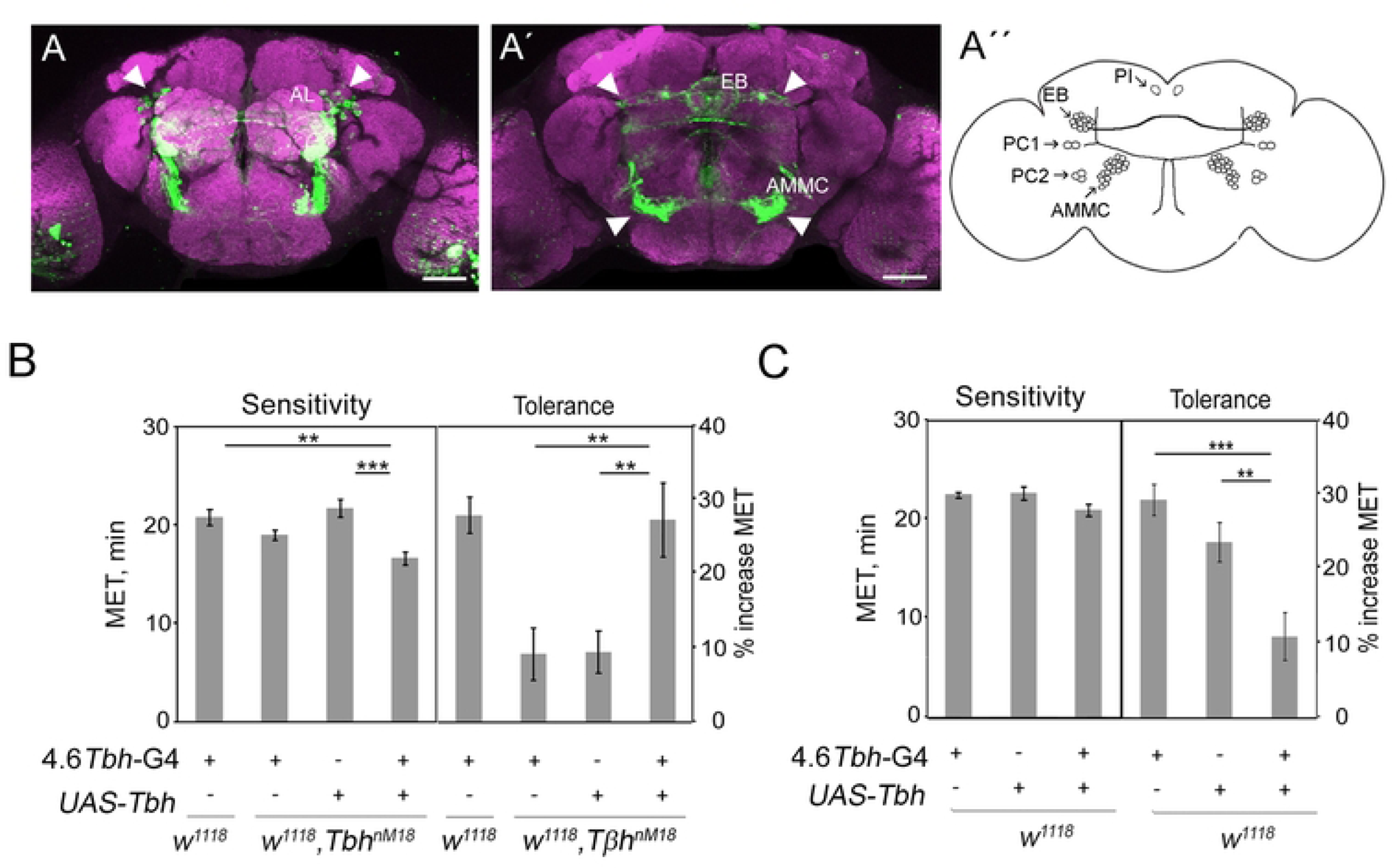
Tbh expression under the control of the 4.6-*Tbh*-Gal4 restored the reduced tolerance to control levels. (A) The Gal4 expression pattern of the *4.6-Tbh*-Gal4 driver is visualized using a *UAS-mCD8::*GFP transgene (in green). The nc82 antibody is used to label Neuropil, (in magenta). Shown are the the Z-projections of the anterior (A) and posterior (À) parts of a confocal stack. Antennal lobes (AL), ellipsoid body (EB), antennal-mechanosensory and motor center (AMMC) are labeled and the white arrowheads indicate soma or projections. Expressing of *Tbh* in a *4.6-Tbh*-Gal4-dependent manner restored the reduced tolerance of *Tbh^nM18^* mutants to control levels (N = 14 - 18). (C) Overexpression of *Tbh* using the *4.6-Tbh*-Gal4 driver significantly reduced ethanol tolerance (N = 7). The ANOVA Tukey-Kramer *post hoc* test was used to determine significance. The asterisks represent \*\**P* < 0.01 and \*\*\**P* < 0.001 and the error bars represent s.e.m.. For data see S2 Table.

### Identification of Tbh-positive cells in the adult brain

We next wanted to examine whether the neurons targeted with the *4.6-Tbh*-Gal4 driver also express Tbh. Therefore, we raised an antibody against the amino acid sequence 112 - 562 of the Tbh isoform Dm-I (Tbh-AB450). The immunoreactivity detected by Tbh-AB450 was still present in *Tbh^nM18^* mutants. To validate the specificity of the antibody serum for Tbh, we compared the immunoreactivity detected by the Tbh antibody to the expression of an endogenous Tbh::GFP fusion protein (Fig 6A and Fig 6B). The *Tbh^fTRG00885.sfGFP-TVPTBF^* insertion line carries a genomic fragment in which the GFP gene has been inserted in frame to the C terminus of the *Tbh* gene (23). Because the C-terminal region is shared by different *Tbh* transcripts, the expression of the Tbh::GFP fusion protein should reflect the expression of different Tbh isoforms. Expression of the Tbh::GFP fusion protein was detected in a large number of cells. In some of the cells, its overlapped with immunoreactivity detected by the antibody. For example, in the anterior part of the brain, the Tbh::GFP expression overlaps with the lateral anterior cells (La) and in the posterior part cells of the posterior cluster (PC1) with Tbh immunoreactivity (Fig 6A and Fig 6B). The results show that the Tbh-AB450 antibody recognizes Tbh and also demonstrate further that more than one Tbh isoform exists.

**Figure 6:**
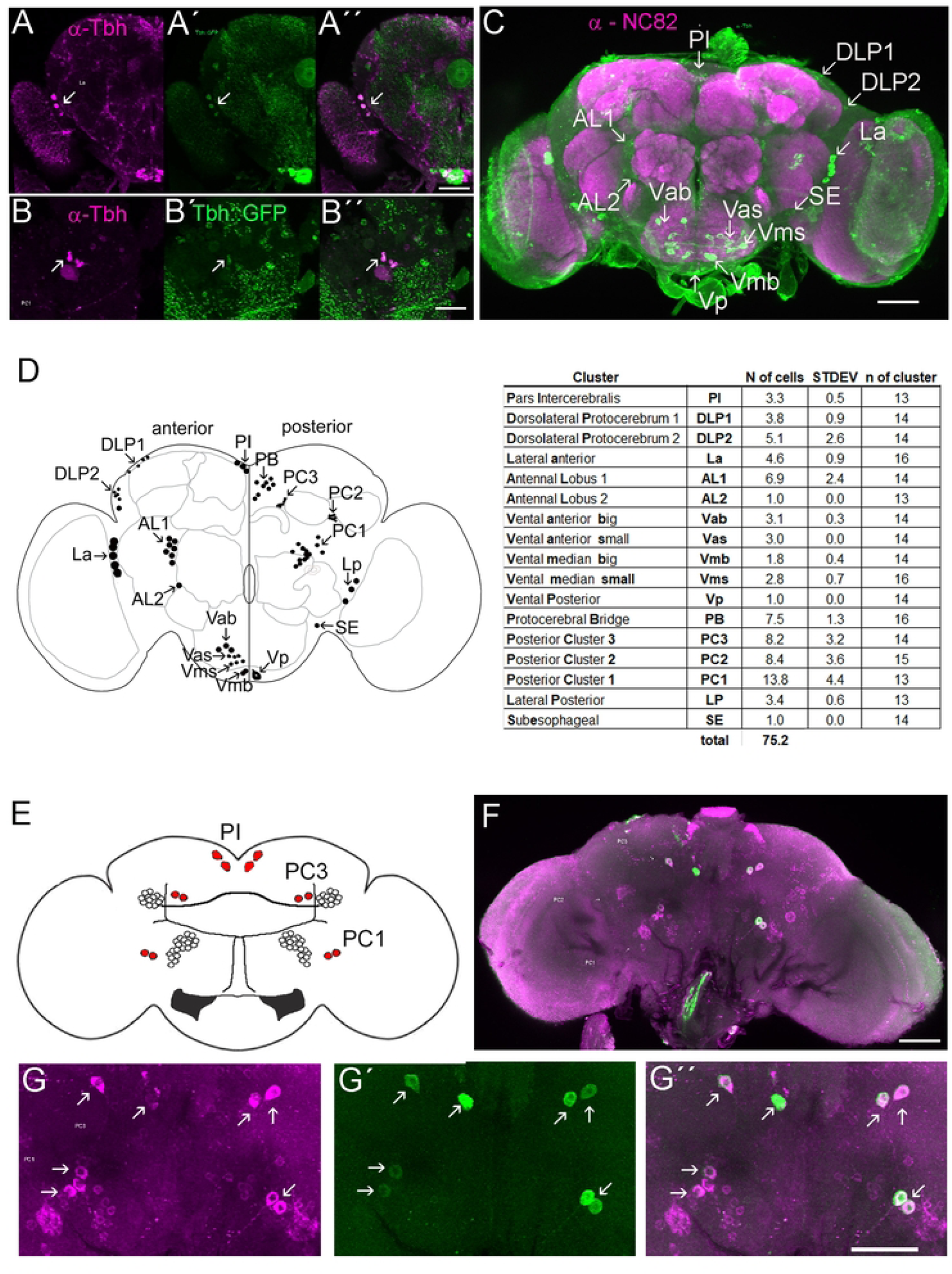
Tbh expression in the adult brain. Tbh::GFP expression of the *Tbh^fTRG00885.sfGFP-TVPTBF^* insertion line in the anterior part of the brain (A) and posterior part (B) is shown in green and overlaps with Tbh (in magenta) in some cells (in white). (C) In the adult brain, the immunoreactivity detected by the Tbh antibody serum is marked in green and the neuropil is marked in magenta by the marker NC82. (D) The scheme and table summarize the immunoreactivity detected by the Tbh antibody serum. (E) The schematic shows a summary of the Gal4 expression domain of the *4.6-Tbh*-Gal4 driver. Cells expressing GFP only are indicated by open circles, and cells expressing GFP and Tbh are indicated as red circles. (F) The Gal4 expression domain of the *4.6-Tbh*-Gal4 driver, visualized with the *UAS-mCD8::*GFP, co-localize with Tbh (in magenta) and GFP (in green) in posterior cells of the brain. The posterior region including the PC1 and PC3 clusters is shown in higher magnification in (G). The white arrow indicates cells co-expressing Tbh and GFP. Scale bars are 50 µm.

We next analyzed the Tbh expression in the adult brain in more detail (Fig 6C to Fig 6D). In the anterior part of the brain, Tbh-positive cell soma are found in the pars intercerebralis, dorsolateral protocerebrum in two clusters, lateral to the optic lobes, adjacent to the antennal lobes in two clusters and in the ventral region of the esophagus (Fig 6C). In the posterior part, Tbh-positive reactivity is found in the soma of cells in three clusters, near the protocerebral bridge, near the lobula plate, and in the lateral region and ventral region of the esophagus. Overall, in the adult brain, the Tbh immunoreactivity was found in 17 clusters containing approximately 75 cells (Fig 6D).

To investigate whether the neurons targeted with the *4.6-Tbh*-Gal4 driver also expressed Tbh, we compared its expression pattern using a *UAS*-mCD8::GFP transgene to the immunoreactivity detected by the Tbh-AB450 antibody (Fig 6E to Fig 6G). In the posterior brain, the Tbh expression is present in 4 cells in the pars intercerebralis (PI) and in 8 cells of the PC1, which overlaps with the GFP expression pattern of the 4.6-Tbh-Gal4 driver. Taken together, expression analysis of the 4.6-Tbh-Gal4 driver and behavioral experiments uncovered the cells in which Tbh is required for the regulation of ethanol tolerance.

### Different Tbh isoforms are expressed in different cells in the larval brain

To analyze whether different Tbh isoforms are expressed in the same or different neurons, we examined the expression of Tbh in the larval central nervous system (CNS) of the 3^rd^ instar with two different Tbh antibodies. We used the Tbh-AB450 that were raised against the amino acid sequence 112 - 562 of the Tbh isoform Dm-I and the Tbh-AB670 that were raised against the amino acid sequence 1 – 670 of the Tbh isoform Dm-I (Tbh-AB-670; (6).

We first wanted to confirm that both antibodies recognize Tbh proteins. Therefore, we compared the immunoreactivity detected by each Tbh antibodies to the expression of an endogenous Tbh::GFP fusion protein using the *Tbh^fTRG00885.sfGFP-TVPTBF^* insertion line (Fig 7A and Fig 7G). Expression of the Tbh::GFP fusion protein is detected in a large number of neurons and overlap in the subesophageal region with Tbh immunoreactivity detected by the antibody Tbh-AB450 (Fig 7A and Fig 7A′′). The antibody Tbh-AB670 also recognized GFP-positive cells in the subesophageal region and cells along the ventral midline (Fig 7G and Fig 7G′′). Both antibodies labeled cells that were only recognized by each antibody individually but were not GFP-positive. In addition, there were also GFP-positive neurons that were not recognized by either antibody. To analyze whether both antibodies recognized the same neurons in the subesophageal region, we compared the immunoreactivity detected by both antibodies against the expression domain of the *dTdc2*-Gal4 driver, which was visualized using the *UAS*-mCD8::GFP transgene (Fig 7B to Fig 7F and Fig 7H to Fig 7L). In the larval brain, the antibody Tbh-AB 450 detected Tbh immunoreactivity only in cells not targeted by the *dTdc2*-Gal4 driver (Fig 7B to Fig 7F), mainly in the soma and punctate structures in both brain hemispheres (Fig 7C), in the subesophageal region (Fig 7D), and the abdominal segments (Fig 7F). In contrast and similar to the adult nervous system, the immunoreactivity detected by Tbh-AB670 antibody(6) partially overlapped with the expression domain of the *dTdc2*-Gal4 driver (13); Fig 7H to Fig 7L). The Tbh-AB670 antibody recognized approximately 48 cells, of which approximately 19 cells are also GFP-positive in the CNS of the larvae (Fig 7H to Fig 7L). Co-localization of Tbh-positive cells that also expressed GFP occurred in 18 out of 20 in the subesophageal medial region (SM) (Fig 7I), and in one cell per hemi-segment in the thoracic region of the ventral nerve cord (VNC) (Fig 7K). None of the Tbh-positive immunoreactivities appear to co-localize with GFP in the brain lobes or in the abdominal region (Fig 7H and Fig 7H′′). However, punctate immunoreactivity can be seen in the abdominal region, which has phenotypic similarities to synaptic boutons but does not overlap with GFP expression (Fig 7L). In the larval and adult CNS, the *dTdc2*-Gal4 driver targets octopaminergic neurons (22); (24). Thus, the Tbh-AB670 recognizes octopaminergic neurons, consistent with the function of Tbh in octopamine synthesis. However, the second antibody Tbh-AB450 recognizes a Tbh-isoform that is not expressed in octopaminergic neurons, which suggests that Tbh function is decoupled from octopaminergic neurons.

**Figure 7:**
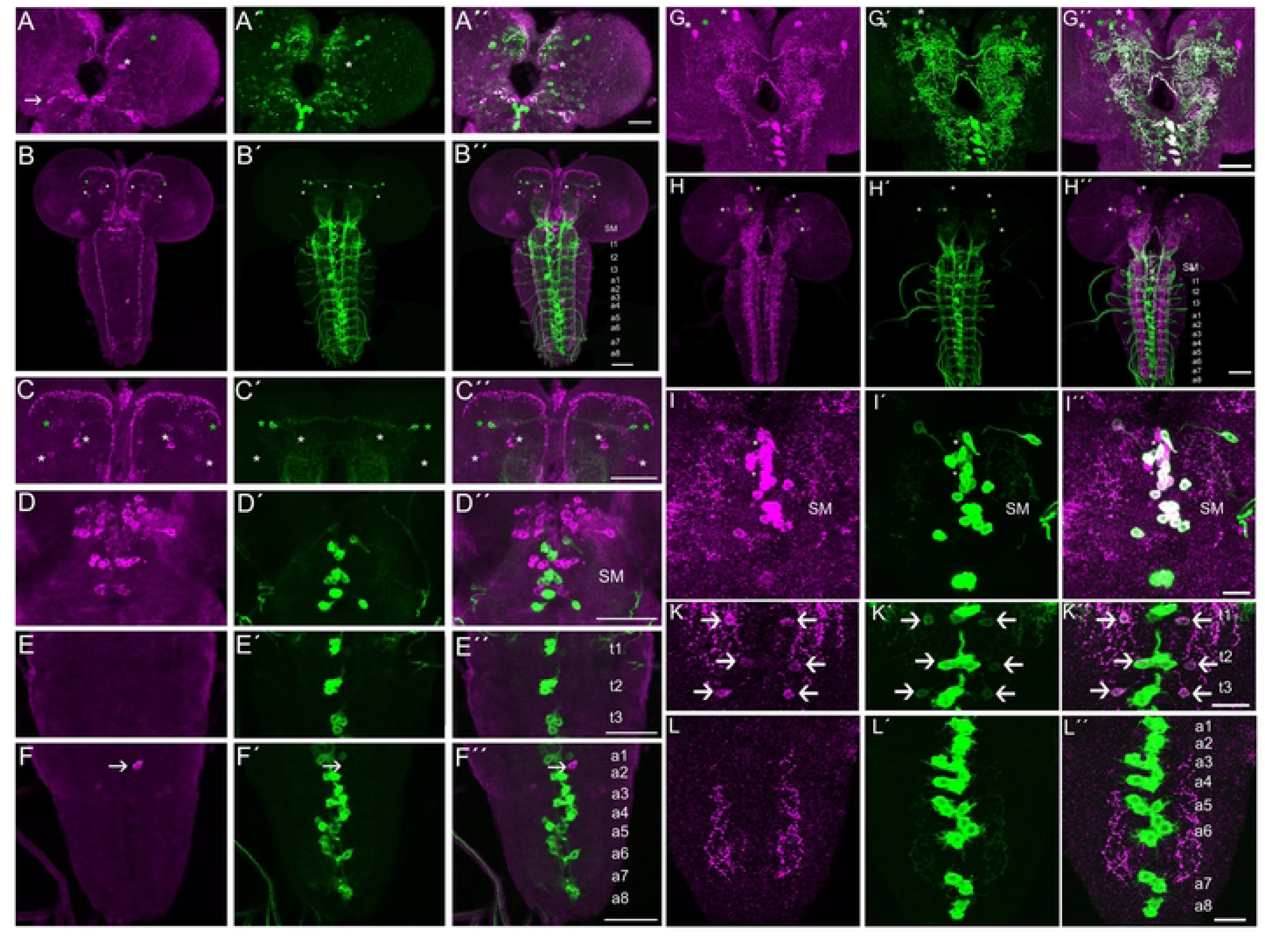
Tbh isoform expressions in the larval brain. In the CNS of the 3^rd^ larval instar, Tbh immunoreactivity (in magenta) is detected with the anti-Tbh antibody serum raised against amino acid 111 - 562 (A to F) and with the anti-Tbh antibody serum raised against amino acid 1 - 670 (G to L; Zhou et al., 2008). In (A) and (G), the Tbh::GFP fusion protein expressed by the *Tbh^fTRG00885.sfGFP-TVPTBF^* insertion line is marked in green. In (B to F and H to L), the expression of the *dTdc2*-Gal4 targeted neurons is visualized by the *UAS*-mCD8::GFP transgene (in green).The overlap between GFP and Tbh is marked in white. The white stars mark cells that are only Tbh-positive, green stars mark cells that express only GFP, and the arrows mark cells that express Tbh and GFP. In (C to F and I to L) a higher magnification of the larval brain is shown: in (C) the brain hemispheres, in (D and I) the subesophageal medial region (SM), in (E and K) the thoracic region t1 to t3 and in (F and L) the abdominal region a1 to a8. The scale bar represents 50 µm.

### One Tbh isoform is expressed in noradrenergic neurons

The expression pattern recognized by the Tbh-AB450 in the subesophageal medial region is similar to the expression pattern of neurons expressing the neuropeptide Hugin (Hug) (25). This prompted us to analyze whether the Tbh immunoreactivity co-localize with the expression pattern of the *HuginS1-*Gal4 driver (Fig 8A to Fig 8B). In the larval brain hemispheres, GFP expression is colocalized with Tbh immunoreactivity in punctate structures and in the VNC with projections of HUG^VNC^ neurons ((26); Fig 8C and Fig 8D). In the subesophageal region, the expression pattern of the *HuginS1*-Gal4 driver strongly overlaps with Tbh immunoreactivity in somata of the Hug^RG^, Hug^PC^, Hug^VNC^ and Hug^PH^ (Fig 8E).

**Figure 8:**
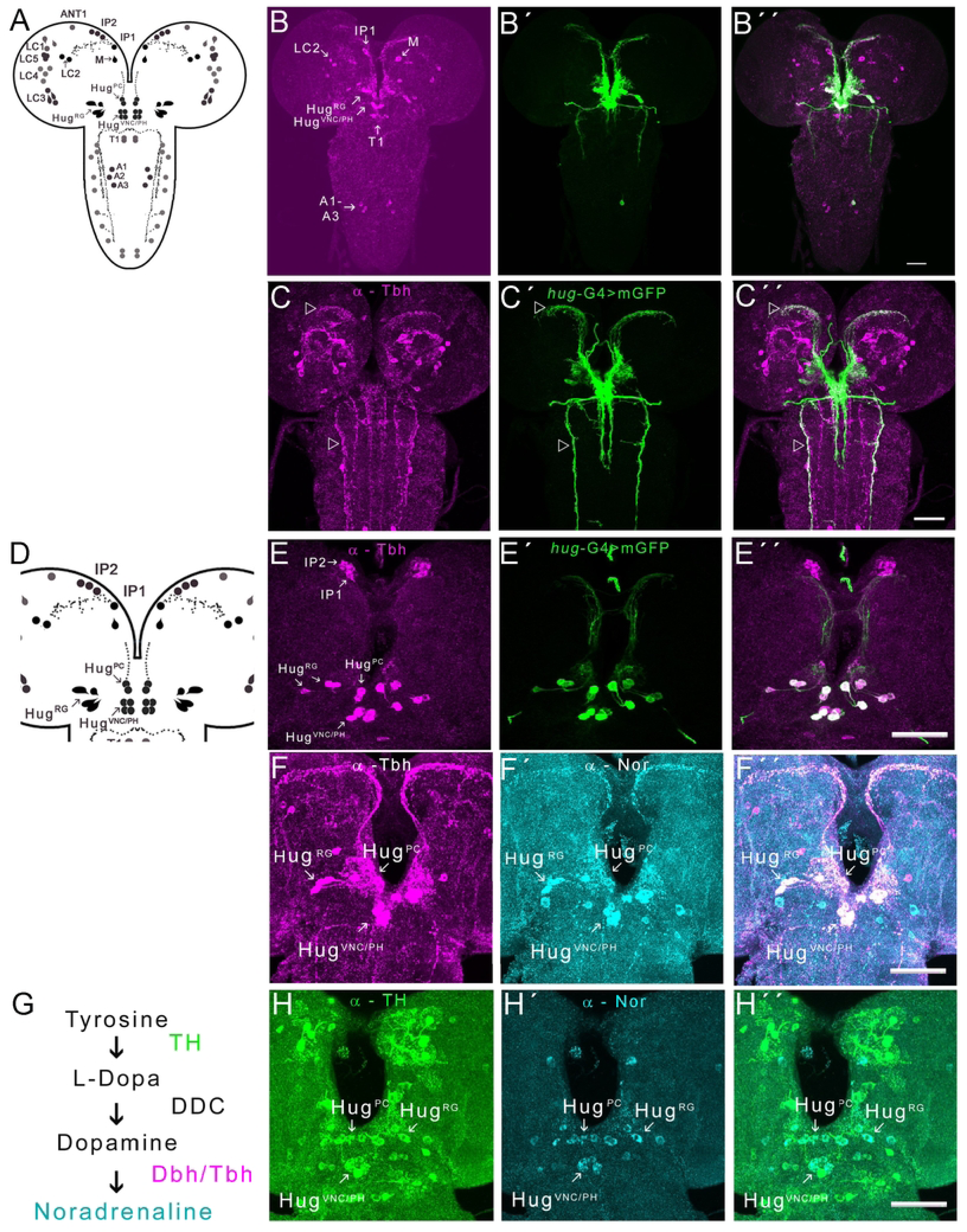
Tbh is expressed in noradrenergic neurons. (A) Schematic overview of Tbh-positive neurons in larval CNS and in (D) a magnification of the subesophageal region for better references. In (B to F) the Tbh-positive immunoreactivity detected by anti-Tbh antibody serum raised against amino acid 111 – 562 is marked in magenta and in (B to E), the expression domain of the *HuginS1*-Gal4 driver is visualized by *UAS*-mCD8::GFP transgene in green. In (F′ and H′) neurons that are noradrenaline-positive are labeled in cyan and in (H) neurons that express TH in green. Neurons that co-express Tbh and GFP or noradrenaline appear white and neurons that express TH and noradrenaline appear light blue. In (G) the noradrenaline synthesis is shown. The arrow heads in (C) indicate areas of synaptic varicosities and projections of neurons co-expressing Tbh and GFP. The arrows mark cells that are Tbh or Tbh and GFP positive. The scale bar represents 50 µm.

Since Tbh shares similarities with the mammalian Dbh and Dbh is required for the conversion of dopamine to noradrenaline (1), (3), we reasoned that the Tbh-AB450 may recognize noradrenergic neurons. To test this, we performed colocalization studies with Tbh and noradrenaline (Fig 8F). In addition to synaptic varicosities in the area of the pars intercerebralis, the somata of the Hug^RG^, Hug^PC^, Hug^VNC^ and Hug^PH^ are Tbh and noradrenaline-positive. To further test whether these neurons are able of producing the precursor for noradrenaline synthesis, dopamine, we analyzed whether these neurons also express the rate-limiting enzyme of the dopamine synthesis, Tyrosine hydroxylase (TH), which indeed was the case (Fig 8H). In summary, the Tbh serum recognizes neurons that express TH, the rate-limiting enzyme of dopamine synthesis, and are noradrenergic neurons (Fig 8G). However, Tbh-AB450 does not detect octopaminergic dTdc2-positive neurons (Fig 7). This suggests that a second Tbh isoform exists that links Tbh function to noradrenaline synthesis.

## Discussion

The *Tbh* gene encodes at least three different isoforms that differ in their size, post-translational modification sites and functional domains. Two of the identified transcripts differ in their 5’-UTR and have the same coding sequences, suggesting that the translation is differentially regulated. At protein level, the different isoforms differ in the number of six putative protein kinase C phosphorylation sites. For example, the isoform Tbh^PC^ contains six putative protein kinase C phosphorylation sites, while Tbh^PD^ has only two sites. The differences may lead to changes in Tbh enzyme expression and activity. Changes of Tbh activity have been observed during development and upon exposure to stress. In the moth *manduca sexta,* the activity of Tbh increases during adult development (27), and in the cockroach *Periplaneta Americana,* the activity of Tbh increases shortly after mechanical stressful stimulation (16).

The newly generated *Tbh^Del3^* mutants exhibit similar locomotor and ethanol tolerance defects as the *Tbh^nM18^* mutants. The reduced locomotion of *Tbh* mutants is due to a lack in adaptation in motor performance rather than defects in behavioral execution, since providing a strong environmental stimulus can increase locomotion. Similarly, they develop reduced tolerance after a previous ethanol exposure but eventually reach the same level after multiple ethanol doses or with chronic ethanol administration (10); (15). The development of tolerance is an adaptation to previous experience. The observation that the behavioral defects in *Tbh* mutants can be modulated by external stimuli or changes of internal conditions supports that the octopaminergic system regulates the behavioral outcome (12).

The inability to properly regulate the development of ethanol tolerance can be traced to a group of eight neurons in the posterior part of the adult brain. One brain region involved in the regulation of ethanol tolerance is the central complex (10); (28). A pair of F1 neuron within the central complex mediates Dunce-dependent ethanol tolerance, as well as experiences dependent changes in locomotion (29); (30). Whether and how the Tbh positive neurons located in the posterior part of the brain are connected to the central complex remains to be seen.

*Tbh^Del3^* mutants differ from *Tbh^nM18^* in the regulation of cellular stress responses. The *Tbh^Del3^* mutants show a reduced response to heat induced ethanol resistance while the *Tbh^nM18^* mutants show a normal response. A stress dependence of Tbh function was also demonstrated indirectly by changes in octopamine levels. In *Drosophila virilis,* a close relative of *Drosophila melanogaster,* heat exposure increases octopamine levels (31). In the locust *Schistocerca gregaria,* mechanical stress leads to the conversion of tyramine into octopamine resulting in an elevation of octopamine levels (32). These results as well as the phenotypic differences of *Tbh* mutants are consistent with the assumption that a stress-inducible and a non-stress inducible octopamine synthesis exist. These differences could be regulated by different *Tbh* transcripts.

Evidence for functional diversity of Tbh isoforms comes from the observation that different Tbh isoforms are expressed in different cells. The expression of one Tbh isoform is mainly detected in tyraminergic/octopaminergic neurons, which link Tbh function to octopamine synthesis. These neurons are consistently octopamine-positive (22). The expression of Tbh is also functionally linked to octopamine synthesis, as *Tbh* mutants do not have detectable octopamine levels (1) and behaviors can be restored by adding octopamine. For example, the loss of ethanol preference in *Tbh^nM18^* mutants can be reversed by feeding octopamine (12).

A second isoform is not expressed in tyraminergic/octopaminergic neurons, but is detected in neuropeptide Hugin-positive neurons. The question arises whether the synthesis of octopamine must necessarily be linked to tyramine synthesis in the same cells. Tyramine can be taken up by food that already contains tyramine or from neighboring cells. For example, a mixture of yeast and bacteria can produce tyramine (33) and glial cells can take up tyramine (34). However, a putative other substrate for Tbh is dopamine. The Tbh shares 39% of its identity with the mammalian Dopamine beta-hydroxylase, which is required for the conversion from dopamine into noradrenaline (1).

The rate-limiting enzyme of dopamine synthesis, Tyrosine hydroxylase, is expressed in the same cells as the Tbh isoform. Furthermore, an antibody to noradrenaline with high affinity for noradrenaline but very low cross-reactivity for octopamine recognizes noradrenaline in the same neurons. The possibility that noradrenaline is indeed synthesized in the fly brain is further supported by the detection of noradrenaline by mass spectrometry (35). It is possible that noradrenaline is synthesized but does not function because a receptor is missing. Most of the *Drosophila melanogaster* G-protein-coupled octopamine or tyramine receptors have not yet been studied for their response to noradrenaline. However, the tyramine receptor 3 responds to noradrenaline with internalization and increased cAMP levels, both indicators for functionality (36), making the receptor a good candidate for mediating the effects of noradrenaline. Therefore, it is likely that noradrenaline is functional in *Drosophila melanogaster*.

In summary, Tbh function can be controlled on transcriptional, translational and on posttranslational levels. These mechanisms allow for temporary control of Tbh functions in response to changes induced by for example environmental stressors such as heat. They can also control functional differences in neurotransmitter synthesis.

## Material and Methods

### Fly Stocks

The following lines were used: *Tbh^nM18^* (1), *dTdc2*-Gal4 (37), NP7088-Gal4 (22), *6.2*-*Tbh*-Gal4 (13), *hs-Tbh* (4), *UAS*-*Tbh* (13), *UAS*-mCD8::GFP (38), and *Tbh^fTRG00885.sfGFP-TVPTBF^* (VDRC#318242). The lines *XP^d10000^* and *XP^d01344^* were obtained from Exelixis at Harvard Medical School. Flies carrying transposable elements were backcrossed to the laboratory’s *w^1118^* background for at least five generations. Flies were reared on ethanol-free s medium with a 12-12 h light-dark cycle at 25°C and 60% humidity. For behavioral experiments, three-to five day-old-male flies that had recovered from CO_2_ sedation for 36 h were used.

### Generation of the*Tbh^Del3^* allele

The FLP/FRT recombination was performed as previously described (18). The *XP^d10000^* and *XP^d01344^* lines were used as parental FRT sites carrying P-element insertion lines (obtained from the Exelixis Collection at Harvard Medical School). In transheterozygous flies carrying both XP lines, recombination was induced using a heat shock-inducible Flipase, resulting in a deletion of the sequence between the two FRT sites of the transposons. In order to generate a deletion, the following crosses were set up. In detail, 35 homozygous *XP^d01344^* female virgins were mated with 15 male flies carrying the *hs*-FLP transgene on the third chromosome (*w^1118^*; MKRS, *hs*-FLP/TM6B, Tb^1^ - BDSC #279). The crosses were set up 30 times. Subsequently, 45 crosses with 15 males of the F1 generation were mated with 35 *XP^d10000^/FM7* virgin females. After three days, the parental flies were removed and the larvae were exposed to heat shocked in a 37 °C water bath for one h on four consecutive days. In the next generation, 35 virgin females of the F2 generation were mated to 15 males carrying the first chromosomal balancer Binsinscy (BDSC #7755). In the F3 generation, 350 individual male flies were mated to three to five *Tbh^nM18^*/FM7 virgins to establish a stock and to select for female sterility. Next, males of the newly generated stocks, which were also female-sterile, were screened for the genomic deletion using PCR analysis.

First, the 5’ and 3’ flanking sequences associated with XP insertion lines were confirmed. To map the *XP^d01344^* insertion the following primers were used: 5’– TGGCACACACTTACGGGTTA-3’ and 5’-GGGAAACGCCTGGTATCTTT-3’ and for the *XP^d01000^* insertion: 5’-GTGCAAAGTGCTCACGCTTA-3’ and 5’–GGGAAACGCCTGGTATCTTT-3’. The deletion contains only fragments of the *XP^d01344^* insertion. The following primers were used to map the truncated *XP^d10000^* P-element insertion: 5’-TTAGCTGCACATCGTCGAAC3’ and 5’-AGCCGGATGACATTATCTGC-3’. The deletion was confirmed using the following primers: 5’-ATTCCGCTGCAGCTGAGCAG-3’ and 5’-GGACTGACACTCACGGAGACA-3’. Of approximately 40 female sterile lines, 5 lines carried the deletion. Before use, the newly generated *Tbh^Del3^* mutant and the parental XP-element insertion lines were backcrossed with *w^1118^* for five generations to isogenize the background.

### Generation of the *4.6*-*Tbh*-Gal4 driver

The 4.6 kb promoter fragment of the *Tbh* gene ranging from - 877 to + 3718 was amplified using the primers 5’-TGGCACACACTTACGGGTTA-3’ and 5’-GGTGGTGATGGAGTCGCC-3’. The fragment was cloned into the p221-4 GAL4 vector via the pCRII vector (LifeTechnologies) using the BamHI and NotI restriction sites. The construct was injected into *w^1118^* embryos to generate transgenic lines. We used random insertions to generate lines with different Gal4 expression patterns and different chromosomal insertions. Insertion #6.5 on the second chromosome was chosen for the experiments.

### Behavioral assays

Sensitivity and tolerance to ethanol were tested as previously described (19); (10). Briefly, 120 flies were introduced into a column filled with ethanol vapor. The average time a population spent in the column, defined as *mean elution time* (MET), was determined. The MET1 reflects the initial sensitivity to the effect of ethanol on postural control, while the MET2 refers to the second exposure to ethanol (19); (10). After the first exposure, flies were collected and reinserted into the columns after a recovery period of four h. The tolerance was calculated as follows: (MET2 – MET1) / MET1*100. To determine the effect of a heat shock on ethanol-induced behavior, 100 flies in an empty vial were incubated in a water bath at 37 °C for 30 min, followed by a 3.5-h recovery at 25 °C. The flies were then exposed to ethanol in the inebriometer (MET+hs) Tolerance was calculated in comparison to non-heat-treated flies (MET-hs).

### Statistics

The Shapiro-Wilk test (significance level *P* < 0.05) followed by a QQ-Plot chart analysis were used to analyze whether the data were normal distributed. For normal distributed data, significance was determined using the Student’s paired *t* - test or using the one-way ANOVA combined with the Tukey-Kramer *post-hoc* HSD test. For nonparametric data, the Mann-Whitney U test for differences between two groups and for more than two groups the Kruskal-Wallis test with post hoc Duenn analysis and Bonferroni correction were used. The statistical data analysis was performed using statskingdom (https://www.statskingdom.com). The errors are indicated as standard error of the mean (s.e.m.).

### Quantitative Real-Time PCR

Total RNA was isolated from fly heads by acid guanidinium thiocyanate-phenol-chloroform extraction using TRIzol® reagent (Invitrogen). For cDNA synthesis, we used oligo-dT primers and the SuperScript^TM^ II reverse transcriptase. Control primers were selected using the NormFinder software (39). The level of the *Tbh* transcripts was normalized to the transcription of the reference gene *RpLP0* (5’-CAGCGTGGAAGGCTCAGTA-3’ and 5’-CAGGCTGGTACGGATGTTCT-3’). Data are shown as fold changes of *Tbh* transcription relative to the reference primers. The primers used to detect the *Tbh^5-6^* region were 5’-ATCCGTACGTTCGACTGGAG-3’ and 5’-TCGACATCTTGATGCGAAAG-3’. The primers used to detect the induction of *hsp70* transcription were 5’-GGCATATCTGGGCGAGAGCATC-3’ and 5’-CTTGAACTCGTCCGCCAGATGAG and for *Rap2l* 5’-ACTTCCGTGCATTACGTGCG-3’ and 5’-CCGACCCGAGCACAACAACT-3.

### Identification of *Tbh* transcripts

To analyze *Tbh* transcripts, we isolated total RNA of three-to five-day-old adult *w^1118^* fly heads using TRIzol^®^ reagent (Invitrogen). The cDNA was generated using oligo-dT primers and the SuperScript^TM^ II reverse transcriptase. To identify additional sequences at the 5’ end, we analyzed the sequences of *Tbh* EST clones. The cDNA clone EY198604.1 contained additional sequence at the 5’ prime end. The sequence was confirmed using the 5’ prime end specific 5’-ACGCGCTTTCCACTTGTTCG-3’ primer. To identify additional transcripts, we used primers that recognize sequences in the 5’ prime region combined with primers that recognize sequences in the 3’ prime region to amplify splice variants (the 5’end primer 5’-ACGCGCTTTCCACTTGTTCG-3’ combined with the 3’end primer 5’-GCTTTCGCTTGGTTTTTGTT-3’). The exon 3 was identified by analyzing putative open reading frames in intron three and designing primers against the sequence. To validate the new exon 3, polyA selected RNA was isolated from total RNA of 1000 heads of one-to two-day-old flies using the MicroPoly(A)Purist™ Kit (Ambion). After cDNA synthesis using oligo-dT primers, nested PCR were performed using the LongAmp^®^ *Taq* DNA Polymerase. For the first synthesis, the following primers were used: 5’-CGAGTGCGATGCATCAAGTG-3’ and 5’-CCATTCGATGCTCCGGTAAT-3’ and for the second synthesis: 5’-CCCAAAAGGGTCGTTCTGTC-3’ and 5’-CAGTTTGGTGGCCGGATAG-3’. The isolated fragment of 1.5 kb was subcloned and sequenced.

### Generation of Tbh antibody serum

The pBluescriptSK-*Tbh* vector containing the *Tbh* cDNA (Monastirioti et al., 1996) was cut with SalI and XhoI. The SalI/XhoI cDNA fragment coding for the amino acid 112 - 562 of the Tbh enzyme was ligated into the pET28b vector. The vector was transformed into BL21 cells and IPTG was used to induce the expression of the HIS-tag Tbh fusion protein. The fusion protein was purified using Ni-NTA Resin Spin Columns (QIAGEN). Eurogentec (Belgium) carried out the immunization in rabbits. The first bleed (1:500) was used for immunohistochemistry.

### Immunohistochemistry

Brains of three- to eight-day-old male flies were used for immunohistochemistry. The dissection, fixation and staining of the nervous system were performed as previously described (13). To detect Tbh immunoreactivity, fly brains were fixed for 2.5 h at 4 °C and blocked with 2.5% BSA and 5% NGS for one h. The brains were incubated with the primary antibody for two nights at 4 °C and the secondary antibody for one night at 4 °C. The following antibodies were used: rabbit anti-Tbh (1:500; we thank Yi Rao for the generous gift of antibody serum); mouse anti-TH (1:200; ImmunoStar ID: 22941); rabbit anti-noradrenaline (1:500; Chemicon® AB120); chicken anti-noradrenaline (1:500; Antigenes); mouse anti-nc82 (1:20, DSHB), chicken anti-GFP (1:1000; Invitrogen,), rabbit anti-GFP (1:1000, Invitrogen), AlexaFluor 488 goat anti-chicken IgG (1:1000; Invitrogen), AlexaFlour 488 goat anti-rabbit (1:1000, Invitrogen); AlexaFlour 546 goat anti-mouse (1:500, Invitrogen), Cy3 goat anti-rabbit IgG (Jackson Immunoresearch, 1:200). Optical sections of one µm were generated using a Zeiss LSM 510 laser scanning microscope. The data analysis of the confocal stacks was performed using Fiji (ImageJ).

Strains and plasmids are available upon request. The authors certify that all data necessary to confirm the article’s conclusions are included in the article, Figs and supporting information.

### Contributions

Conceptualization: MR, SH, HS; Formal analysis: MR, SH, GC, AD, CF, TK, HS; Funding acquisition: HS; Investigation: MR, SH, NS, OC, GC, AD, CF, TK, SP; Resources: MR, SH, OC; Visualization: MR, HS; Writing: HS.

## Acknowledgement

We thank Alina Bouwer, Sebastian Zetzsche and Evelin Fahle for excellent technical assistance.

## Supporting information

S1 Text: Sequence of *Tbh* transcripts and proteins. S2 Table: Data for behavioral analyses.

## References

1. Monastirioti M, Linn CE, Jr., White K. Characterization of Drosophila tyramine beta-hydroxylase gene and isolation of mutant flies lacking octopamine. The Journal of neuroscience : the official journal of the Society for Neuroscience. 1996;16(12):3900–11.

2. Gray EE, Small SN, McGuirl MA. Expression and characterization of recombinant tyramine beta-monooxygenase from Drosophila: a monomeric copper-containing hydroxylase. Protein expression and purification. 2006;47(1):162–70.

3. Goldstein M, McKereghan MR, Lauber E. The Involvement of Sulfhydryl Sites in Dopamine-Beta-Hydroxylase Activity. Biochim Biophys Acta. 1963;77:161–4.

4. Monastirioti M. Distinct octopamine cell population residing in the CNS abdominal ganglion controls ovulation in Drosophila melanogaster. Dev Biol. 2003;264(1):38–49.

5. Scholz H. Influence of the biogenic amine tyramine on ethanol-induced behaviors in Drosophila. Journal of neurobiology. 2005;63(3):199–214.

6. Zhou C, Rao Y, Rao Y. A subset of octopaminergic neurons are important for Drosophila aggression. Nat Neurosci. 2008;11(9):1059–67.

7. Brembs B, Christiansen F, Pfluger HJ, Duch C. Flight initiation and maintenance deficits in flies with genetically altered biogenic amine levels. The Journal of neuroscience : the official journal of the Society for Neuroscience. 2007;27(41):11122–31.

8. Scheiner R, Steinbach A, Claßen G, Strudthoff N, … Octopamine indirectly affects proboscis extension response habituation in Drosophila melanogaster by controlling sucrose responsiveness. Journal of insect …. 2014.

9. Bainton RJ, Tsai LT, Singh CM, Moore MS, Neckameyer WS, Heberlein U. Dopamine modulates acute responses to cocaine, nicotine and ethanol in Drosophila. Curr Biol. 2000;10(4):187–94.

10. Scholz H, Ramond J, Singh CM, Heberlein U. Functional ethanol tolerance in Drosophila. Neuron. 2000;28(1):261–71.

11. Wolf FW, Rodan AR, Tsai LT, Heberlein U. High-resolution analysis of ethanol-induced locomotor stimulation in Drosophila. The Journal of neuroscience : the official journal of the Society for Neuroscience. 2002;22(24):11035–44.

12. Classen G, Scholz H. Octopamine Shifts the Behavioral Response From Indecision to Approach or Aversion in Drosophila melanogaster. Front Behav Neurosci. 2018;12:131.

13. Schneider A, Ruppert M, Hendrich O, Giang T, Ogueta M, Hampel S, et al. Neuronal basis of innate olfactory attraction to ethanol in Drosophila. PLoS One. 2012;7(12):e52007.

14. Scholz H, Franz M, Heberlein U. The hangover gene defines a stress pathway required for ethanol tolerance development. Nature. 2005;436(7052):845-7.

15. Berger KH, Heberlein U, Moore MS. Rapid and chronic: two distinct forms of ethanol tolerance in Drosophila. Alcohol Clin Exp Res. 2004;28(10):1469–80.

16. Chatel A, Murillo L, Bourdin CM, Quinchard S, Picard D, Legros C. Characterization of tyramine beta-hydroxylase, an enzyme upregulated by stress in Periplaneta americana. J Mol Endocrinol. 2013;50(1):91–102.

17. Aravind L. The WWE domain: a common interaction module in protein ubiquitination and ADP ribosylation. Trends Biochem Sci. 2001;26(5):273–5.

18. Parks AL, Cook KR, Belvin M, Dompe NA, Fawcett R, Huppert K, et al. Systematic generation of high-resolution deletion coverage of the Drosophila melanogaster genome. Nat Genet. 2004;36(3):288–92.

19. Cohan FM, Graf JD. Latitudinal Cline in Drosophila Melanogaster for Knockdown Resistance to Ethanol Fumes and for Rates of Response to Selection for Further Resistance. Evolution. 1985;39(2):278–93.

20. Saraswati S, Fox LE, Soll DR, Wu CF. Tyramine and octopamine have opposite effects on the locomotion of Drosophila larvae. Journal of neurobiology. 2004;58(4):425–41.

21. Fox LE, Soll DR, Wu CF. Coordination and modulation of locomotion pattern generators in Drosophila larvae: effects of altered biogenic amine levels by the tyramine beta hydroxlyase mutation. The Journal of neuroscience : the official journal of the Society for Neuroscience. 2006;26(5):1486–98.

22. Busch S, Selcho M, Ito K, Tanimoto H. A map of octopaminergic neurons in the Drosophila brain. J Comp Neurol. 2009;513(6):643–67.

23. Sarov M, Barz C, Jambor H, Hein MY, Schmied C, Suchold D, et al. A genome-wide resource for the analysis of protein localisation in Drosophila. Elife. 2016;5:e12068.

24. Selcho M, Pauls D, El Jundi B, Stocker RF, Thum AS. The role of octopamine and tyramine in Drosophila larval locomotion. J Comp Neurol. 2012;520(16):3764–85.

25. Melcher C, Pankratz MJ. Candidate gustatory interneurons modulating feeding behavior in the Drosophila brain. PLoS Biol. 2005;3(9):e305.

26. Bader R, Colomb J, Pankratz B, Schrock A, Stocker RF, Pankratz MJ. Genetic dissection of neural circuit anatomy underlying feeding behavior in Drosophila: distinct classes of hugin-expressing neurons. J Comp Neurol. 2007;502(5):848–56.

27. Lehman HK, Murgiuc CM, Hildebrand JG. Characterization and developmental regulation of tyramine-beta-hydroxylase in the CNS of the moth, Manduca sexta. Insect Biochem Mol Biol. 2000;30(5):377–86.

28. Urizar NL, Yang Z, Edenberg HJ, Davis RL. Drosophila homer is required in a small set of neurons including the ellipsoid body for normal ethanol sensitivity and tolerance. The Journal of neuroscience : the official journal of the Society for Neuroscience. 2007;27(17):4541–51.

29. Ruppert M, Franz M, Saratsis A, Velo Escarcena L, Hendrich O, Gooi LM, et al. Hangover Links Nuclear RNA Signaling to cAMP Regulation via the Phosphodiesterase 4d Ortholog dunce. Cell Rep. 2017;18(2):533–44.

30. Martin JR, Ernst R, Heisenberg M. Temporal pattern of locomotor activity in Drosophila melanogaster. J Comp Physiol A. 1999;184(1):73–84.

31. Hirashima A, Sukhanova M, Rauschenbach I. Biogenic amines in Drosophila virilis under stress conditions. Biosci Biotechnol Biochem. 2000;64(12):2625–30.

32. Kononenko NL, Wolfenberg H, Pfluger HJ. Tyramine as an independent transmitter and a precursor of octopamine in the locust central nervous system: an immunocytochemical study. J Comp Neurol. 2009;512(4):433–52.

33. Costantini A, Vaudano E, Del Prete V, Danei M, Garcia-Moruno E. Biogenic amine production by contaminating bacteria found in starter preparations used in winemaking. J Agric Food Chem. 2009;57(22):10664–9.

34. Ryglewski S, Duch C, Altenhein B. Tyramine Actions on Drosophila Flight Behavior Are Affected by a Glial Dehydrogenase/Reductase. Front Syst Neurosci. 2017;11:68.

35. Watson DG, Zhou P, Midgley JM, Milligan CD, Kaiser K. The determination of biogenic amines in four strains of the fruit fly Drosophila melanogaster. J Pharm Biomed Anal. 1993;11(11-12):1145–9.

36. Bayliss A, Roselli G, Evans PD. A comparison of the signalling properties of two tyramine receptors from Drosophila. J Neurochem. 2013;125(1):37–48.

37. Cole SH, Carney GE, McClung CA, Willard SS, Taylor BJ, Hirsh J. Two functional but noncomplementing Drosophila tyrosine decarboxylase genes: distinct roles for neural tyramine and octopamine in female fertility. J Biol Chem. 2005;280(15):14948–55.

38. Lee T, Luo L. Mosaic analysis with a repressible cell marker for studies of gene function in neuronal morphogenesis. Neuron. 1999;22(3):451–61.

39. Andersen CL, Jensen JL, Orntoft TF. Normalization of real-time quantitative reverse transcription-PCR data: a model-based variance estimation approach to identify genes suited for normalization, applied to bladder and colon cancer data sets. Cancer Res. 2004;64(15):5245–50.

